# Sex-Specific Regulation of Stress Susceptibility by the Astrocytic Gene *Htra1*

**DOI:** 10.1101/2024.04.12.588724

**Authors:** Eric M. Parise, Trevonn M. Gyles, Arthur Godino, Omar K. Sial, Caleb J. Browne, Lyonna F. Parise, Angélica Torres-Berrío, Marine Salery, Romain Durand-de Cuttoli, Matthew T. Rivera, Astrid M. Cardona-Acosta, Leanne Holt, Tamara Markovic, Yentl Y. van der Zee, Zachary S. Lorsch, Flurin Cathomas, Juliet B. Garon, Collin Teague, Orna Issler, Peter J. Hamilton, Carlos A. Bolaños-Guzmán, Scott J. Russo, Eric J. Nestler

## Abstract

Major depressive disorder (MDD) is linked to impaired structural and synaptic plasticity in limbic brain regions. Astrocytes, which regulate synapses and are influenced by chronic stress, likely contribute to these changes. We analyzed astrocyte gene profiles in the nucleus accumbens (NAc) of humans with MDD and mice exposed to chronic stress. *Htra1*, which encodes an astrocyte-secreted protease targeting the extracellular matrix (ECM), was significantly downregulated in the NAc of males but upregulated in females in both species. Manipulating *Htra1* in mouse NAc astrocytes bidirectionally controlled stress susceptibility in a sex-specific manner. Such *Htra1*manipulations also altered neuronal signaling and ECM structural integrity in NAc. These findings highlight astroglia and the brain’s ECM as key mediators of sex-specific stress vulnerability, offering new approaches for MDD therapies.

## Main Text

Major depressive disorder (MDD) stands as a formidable public health challenge due to its increasing prevalence and the substantial impact it exerts on individuals and societies worldwide (*1*). The complexity of MDD’s etiology involves a multifaceted interplay of genetic, environmental, and neurobiological factors influencing its onset and progression. The recent COVID-19 pandemic, with its profound social and economic consequences, has further intensified the global burden of depression (*2, 3*). Adding to this complexity, MDD presents a striking sexual dimorphism, with females being 2 to 3 times more susceptible to the disorder and often experiencing more severe symptoms than their male counterparts (*4*). While extensive research has been dedicated to neuronal mechanisms in MDD, there is a burgeoning interest in the role of astrocytes, the predominant glial cells in the brain (*5*). Astrocytes are traditionally recognized for their roles in supporting synaptic structure, function, and neurotransmission (*6*). Recent studies suggest that chronic stress can disrupt astrocyte function, potentially contributing to the striking structural changes observed across limbic reward regions in neuropsychiatric disorders, including MDD (*7*–*9*). Moreover, an accumulating body of evidence connects astrocytic dysfunction to the pathophysiology of MDD (*6, 10, 11*).

Given the profound impact of MDD and the intricacies of its sexual dimorphism, coupled with the emerging importance of astrocytes in understanding its pathophysiology, there is a compelling imperative to comprehensively investigate astrocyte-specific transcriptional profiles in the context of MDD and chronic stress, with a special focus on sex-specificity. In this study, we begin to address this need by analyzing the transcriptional profiles of astrocyte-specific genes within the nucleus accumbens (NAc) – a key brain reward region implicated in depression and stress – obtained from postmortem brain tissue of individuals with MDD and from mice exhibiting some depression-like behavioral abnormalities following exposure to chronic stress.

### *HTRA1* is an astrocyte-specific sexually dimorphic ECM gene influenced by depression and stress in the NAc

We identified high-temperature requirement protein A1 (*HTRA1)*, a secreted serine protease involved in regulating the extracellular matrix (ECM) (*12*), as one of the most significantly differentially regulated astrocyte-specific genes in the NAc of MDD patients and of mice exposed to chronic variable stress (CVS) (Fig. 1A). Upon validation, we discovered that HTRA1 mRNA and protein expression in NAc are sexually dimorphic and oppositely regulated by MDD and CVS in males versus females (Fig. 1B–E). Males display significantly higher baseline expression levels of HTRA1 than females, with a decrease in expression in MDD (Fig. 1B,C) or after CVS (Fig. 1D,E), whereas females exhibit significantly higher expression in both conditions. Although *HTRA1* expression is highly enriched in astrocytes, it is not exclusively expressed in this cell type (*13, 14*). Therefore, we utilized *in situ* hybridization to verify cell-type specificity in CVS-exposed mice and found that CVS-induces *Htra1* mRNA expression in NAc exclusively in astrocytes, with very little expression and no significant differences between conditions in neurons (Fig. 1F,G & Fig. S1) We further confirmed the involvement of *Htra1* in stress responding in male and female mice using a second stress model, chronic social defeat stress (CSDS), and again observed sex-specific effects of CSDS on *Htra1* mRNA expression in NAc (Fig. S2). Taken together, our findings show that *HTRA1* is expressed predominantly in astrocytes of the NAc, and displays sexually dimorphic and opposite regulation in response to chronic stress and MDD, thereby implicating its potential role in mood disorder pathophysiology. Further support for this assertion comes from human studies, where HTRA1 has been identified as a potential blood biomarker for suicidality in male bipolar patients (*15*), and has been found to be differentially expressed in the NAc of individuals with mood disorders who committed suicide compared to those who did not (*16*). However, the causal contribution of HTRA1 to stress susceptibility remains unknown.

**Fig. 1.**
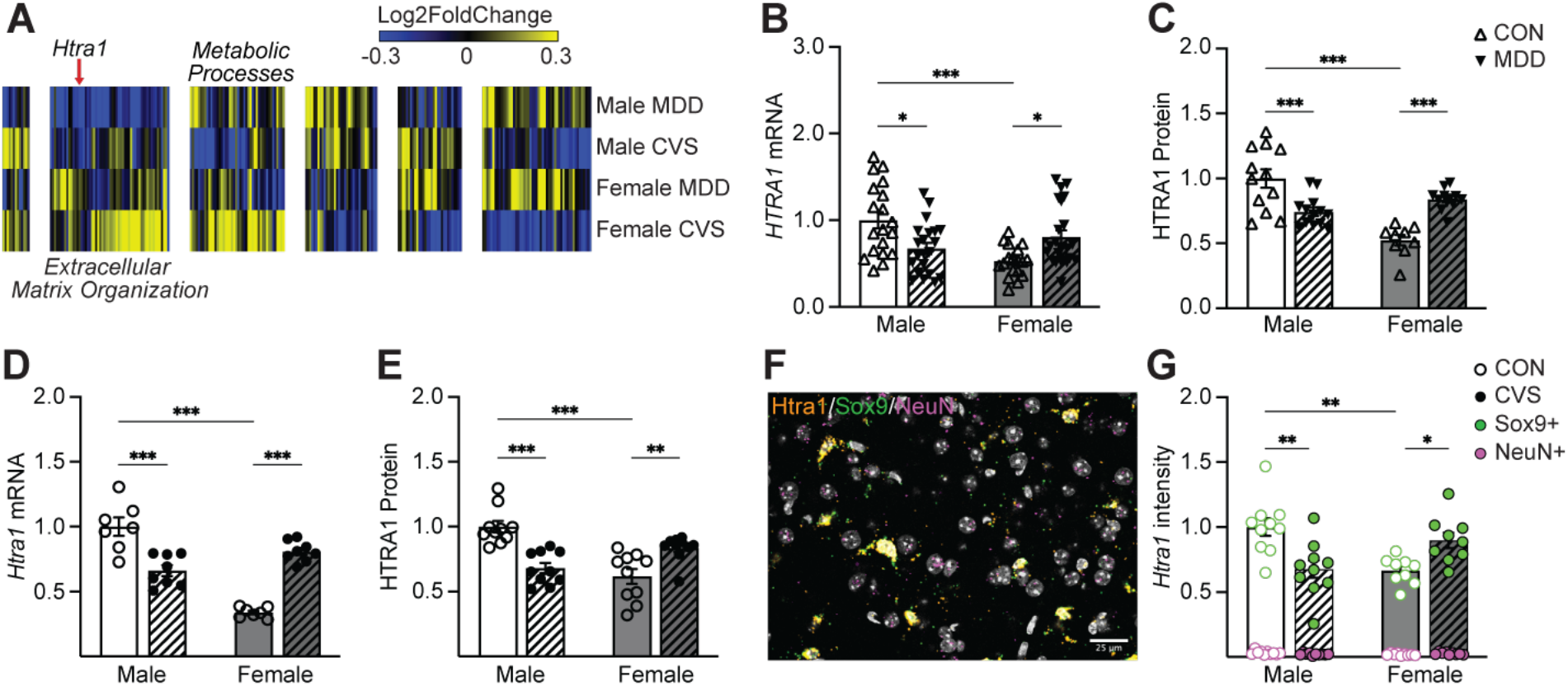
Identification of an astrocyte-specific, sexually dimorphic ECM gene within the NAc that is differentially regulated by depression and stress in a sex-specific manner. (**A**) Union heatmap displaying hierarchical clustering of significantly differentially regulated astrocyte-specific genes in the NAc, identifying high temperature requirement factor A1 (*HTRA1*) as oppositely regulated between males and females both in postmortem human brain tissue from MDD patients and in mice exposed to CVS. Human and mouse RNA-seq data from bulk NAc dissections are from Labonté et al. (*56*). GO analysis reveals enrichment of biological processes for clusters of genes showing distinct patterns across sex, MDD/CVS, and species. (**B**) *Htra1* mRNA (Two-way ANOVA: sex F_1,77_ = 5.387 *p =* 0.0229, MDD F_1,77_ = 0.1001 *p =* 0.7526, sex x MDD F_1,77_ = 17.74 *p* < 0.0001; followed by Tukey’s post-hoc tests) and (**C**) HTRA1 protein (Two-way ANOVA: sex F_1,42_ = 17.01 *p* < 0.0001, MDD F_1,42_ = 0.4025 *p* = 0.5293, sex x MDD F_1,42_ = 38.66 *p* < 0.0001; followed by Tukey’s post-hoc tests) is both sexually dimorphic as well as oppositely regulated in human MDD patients. (**D**) Similarly, in mice exposed to CVS, *Htra1* mRNA (Two-way ANOVA: sex F_1,26_ = 35.11 *p* < 0.0001, stress F_1,26_ = 2.317 *p* = 0.1400, sex x stress F_1,26_ = 86.82 *p* < 0.0001; followed by Tukey’s post-hoc tests) and (**E**) Htra1 protein (Two-way ANOVA: sex F_1,36_ = 6.637 *p* < 0.05, stress F_1,36_ = 1.281 *p* = 0.2651, sex x stress F_1,36_ = 36.99 *p* < 0.0001; followed by Tukey’s post-hoc tests) expression is sexually dimorphic and oppositely regulated between males and females following exposure to CVS. (**F**) Representative confocal image of *Htra1* mRNA expression in neurons (NeuN^+^) vs astrocytes (Sox9^+^) from male (n = 10/group) and female (n = 9–10/group) mice exposed to CVS. (**G**) *Htra1* mRNA expression is significantly higher in astrocytes compared to neurons in the NAc for both sexes, regardless of stress exposure. Importantly, the previously observed sex- and stress-specific patterns of *Htra1 mRNA* expression are observed exclusively in astrocytes (Two-way ANOVA: sex F_1,35_ = 0.9161 *p* = 0.3451, stress F_1,35_ = 0.5966 *p* = 0.4451, sex x stress F_1,35_ = 22.75 *p* < 0.0001; followed by Tukey’s post-hoc tests). All data are normalized to *Htra1* intensity in astrocytes of male controls. \**p* < 0.05; \*\**p* < 0.01; \*\*\**p* < 0.001. Abbreviations: MDD, major depressive disorder; CVS, chronic variable stress; CON, control.

### Viral manipulation of *Htra1* expression selectively in astrocytes of the NAc bidirectionally regulates stress susceptibility in a sex-specific manner

To causally link *Htra1* expression changes to behavioral outcomes, we developed viral constructs to overexpress (OE) or knock down (KD) its expression exclusively in astrocytes using a AAV5-GfaABC1D vector, and confirmed its cell-type specificity as well as OE or KD efficiency (Fig. S3) Next, we bi-directionally manipulated *Htra1* expression in NAc astrocytes in male and female mice and determined their behavioral responses to varying degrees of CVS exposure in a battery of behavioral tests: the splash test (ST), novelty suppressed feeding (NSF), sucrose preference (SP), and open field test (OFT). We found striking sex-specific changes in stress susceptibility upon *Htra1* manipulations in NAc astrocytes (Fig. 2). Astrocytic KD of *Htra1* in the NAc induced behavioral deficits in the ST, NSF, SP, and OFT in male mice following exposure to sub-threshold CVS (ST-CVS) (Fig. 2B–E), whereas in females such *Htra1* KD blocked CVS-induced behavioral deficits in these tasks (Fig. 2F–I). Conversely, astrocytic OE of *Htra1* in the NAc blocked CVS-induced behavioral deficits in the ST, NSF, SP, and OFT in males (Fig. 2J–M), whereas *Htra1* OE in females induced behavioral deficits in these tasks following exposure to ST-CVS (Fig. 2N–Q). These finding establish that bidirectional manipulation of *Htra1* expression in astrocytes within the mouse NAc controls susceptibility to stress in a striking sex-specific manner, with KD inducing stress susceptibility in males but resilience in females, while OE induces resilience in males and susceptibility in females.

**Fig. 2.**
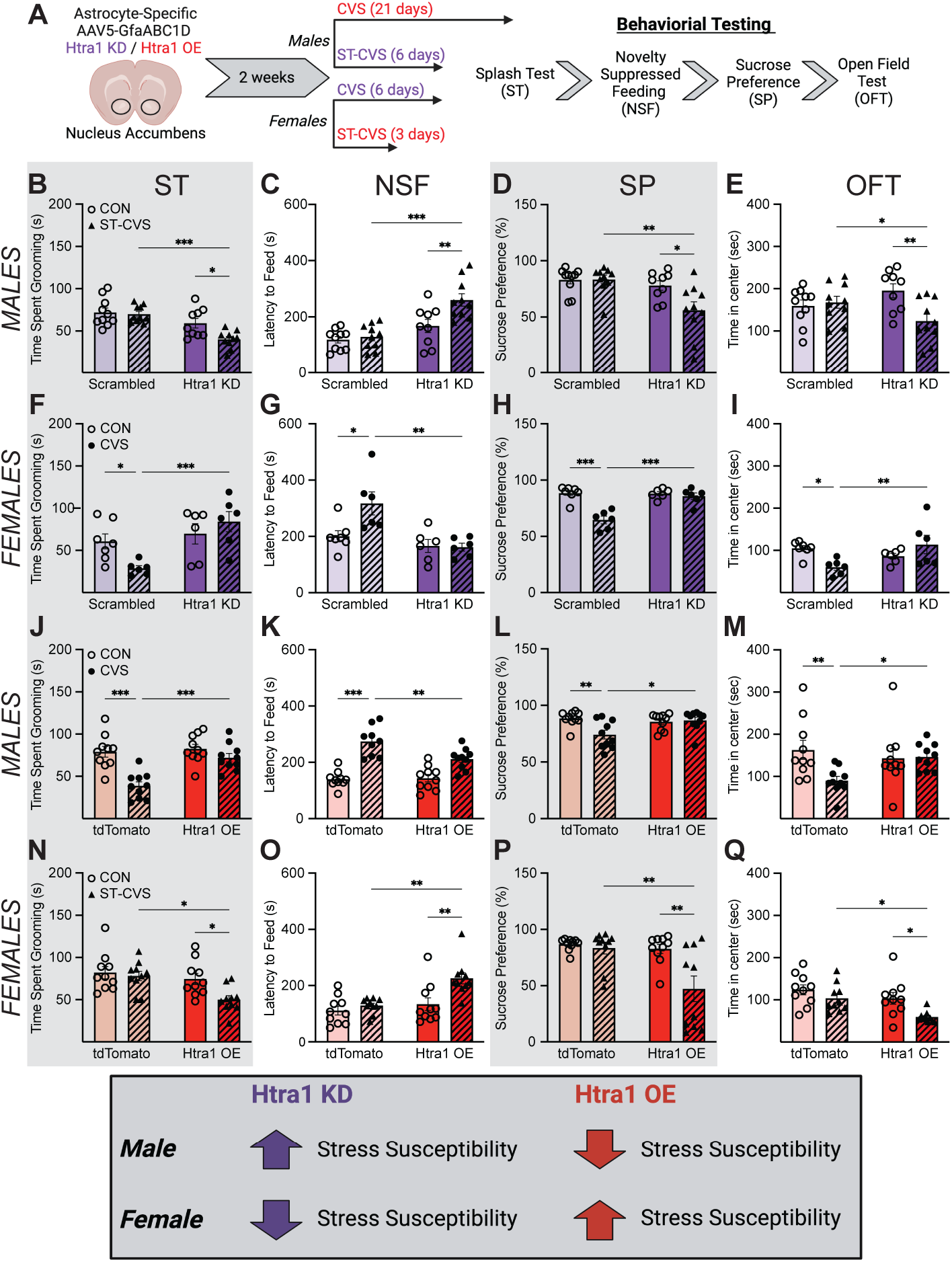
Bidirectional astrocyte-specific manipulation of *Htra1* in the NAc regulates stress susceptibility and resilience in a sex-dependent manner. (**A**) Experimental design and timeline illustrating astrocyte-specific *Htra1* manipulations in NAc of male and female mice. (**B**) In males exposed to ST-CVS, astrocyte-specific *Htra1* KD in the NAc significantly reduced grooming time in the ST (Two-way ANOVA: stress F_1,35_ = 5.743 *p* = 0.0220, virus F_1,35_ = 23.22 *p* < 0.0001, stress x virus F_1,35_ = 3.774 *p* = 0.0601; followed by Tukey’s post-hoc tests), (**C**) increased latency to feed in the NSF task (Two-way ANOVA: stress F_1,35_ = 4.504 *p* = 0.0410, virus F_1,35_ = 10.35 *p* = 0.0028, stress x virus F_1,35_ = 4.875 *p* = 0.0339; followed by Tukey’s post-hoc tests), (**D**) decreased SP (Two-way ANOVA: stress F_1,35_ = 7.983 *p* = 0.0077, virus F_1,35_ = 24.31 *p* < 0.0001, stress x virus F_1,35_ = 5.091 *p* = 0.0304; followed by Tukey’s post-hoc tests), and (**E**) reduced time spent in the center of the OFT (Two-way ANOVA: stress F_1,35_ = 4.353 *p* = 0.0443, virus F_1,35_ = 0.0789 *p* = 0.7804, stress x virus F_1,35_ = 6.987 *p* = 0.0123; followed by Tukey’s post-hoc tests). (**F**) In females, astrocyte-specific KD of *Htra1* blocked the reduction in grooming time (Two-way ANOVA: stress F_1,21_ = 1.408 *p* = 0.2486, virus F_1,21_ = 11.27 *p* = 0.0030, stress x virus F_1,21_ = 7.460 *p* = 0.0125; followed by Tukey’s post-hoc tests), (**G**) prevented a significant increase in latency to feed (Two-way ANOVA: stress F_1,21_ = 4.639 *p* = 0.0430, virus F_1,21_ = 13.30 *p* = 0.0015, stress x virus F_1,21_ = 5.412 *p* = 0.0301; followed by Tukey’s post-hoc tests), (**H**) blocked the decrease in SP (Two-way ANOVA: stress F_1,21_ = 24.14 *p* < 0.0001, virus F_1,21_ = 15.00 *p* = 0.0009, stress x virus F_1,21_ = 16.54 *p* = 0.0006; followed by Tukey’s post-hoc tests), and (**I**) prevented the decrease in time spent in the center of the OFT (Two-way ANOVA: stress F_1,21_ = 0.8824 *p* = 0.3582, virus F_1,21_ = 1.615 *p* = 0.2177, stress x virus F_1,21_ = 11.01 *p* = 0.0033; followed by Tukey’s post-hoc tests) induced by 6 days of CVS (which evokes maximal behavioral responses in female mice (*56, 57*). (**J**) In males, astrocyte-specific OE of *Htra1* blocked the reduction in time spent grooming (Two-way ANOVA: stress F_1,36_ = 22.16 *p* < 0.0001, virus F_1,36_ = 11.64 *p* = 0.0016, stress x virus F_1,36_ = 7.645 *p* = 0.0089; followed by Tukey’s post-hoc tests), (**K**) prevented a significant increase in latency to feed (Two-way ANOVA: stress F_1,35_ = 59.76 *p* < 0.0001, virus F_1,35_ = 4.966 *p* = 0.0324, stress x virus F_1,35_ = 3.774 *p* = 0.0153; followed by Tukey’s post-hoc tests), (**L**) blocked the decrease in SP (Two-way ANOVA: stress F_1,36_ = 5.280 *p* = 0.0275, virus F_1,36_ = 2.919 *p* = 0.0961, stress x virus F_1,36_ = 7.499 *p* = 0.0095; followed by Tukey’s post-hoc tests), and (**M**) prevented the decrease in time spent in the center of the OFT (Two-way ANOVA: stress F_1,36_ = 5.743 *p* = 0.0220, virus F_1,36_ = 23.22 *p* < 0.0001, stress x virus F_1,36_ = 3.774 *p* = 0.0601; followed by Tukey’s post-hoc tests) induced by 21 days of CVS (which is required to induce maximal behavioral responses in male mice (*56, 57*). (**N**) In contrast, in females exposed to ST-CVS, astrocyte-specific *Htra1* OE in NAc significantly reduced time spent grooming in the ST (Two-way ANOVA: stress F_1,36_ = 5.400 *p* = 0.0259, virus F_1,36_ = 8.379 *p* = 0.0064, stress x virus F_1,36_ = 2.794 *p* = 0.1033; followed by Tukey’s post-hoc tests), (**O**) increased latency to feed in the NSF task (Two-way ANOVA: stress F_1,36_ = 9.840 *p* = 0.0034, virus F_1,36_ = 11.67 *p* = 0.0016, stress x virus F_1,36_ = 4.465 *p* = 0.0416; followed by Tukey’s post-hoc tests), (**P**) decreased SP (Two-way ANOVA: stress F_1,36_ = 8.724 *p* = 0.0055, virus F_1,36_ = 9.387 *p* = 0.0041, stress x virus F_1,36_ = 5.862 *p* = 0.0206; followed by Tukey’s post-hoc tests), and (**Q**) decreased time spent in the center of the OFT (Two-way ANOVA: stress F_1,36_ = 8.802 *p* = 0.0053, virus F_1,36_ = 9.041 *p* = 0.0048, stress x virus F_1,36_ = 1.241 *p* = 0.2726; followed by Tukey’s post-hoc tests). \**p* < 0.05; \*\**p* < 0.01; \*\*\**p* < 0.001. Abbreviations: KD, knockdown; OE, overexpression; CON, control; CVS, chronic variable stress; ST-CVS, subthreshold CVS; ST, splash test; NSF, novelty suppressed feeding; SP, sucrose preference; OFT, open field test.

### Viral manipulation of *Htra1* expression in NAc astrocytes modulates neuronal activity in a sex-specific manner

HTRA1 has been implicated in several cellular processes, including neuroinflammation and synaptic plasticity (*17, 18*). Therefore, stress regulation of HTRA1 expression in astrocytes could potentially affect the functioning of neighboring neurons. D1-type and D2-type medium spiny projection neurons (MSNs) constitute ∼95% of all neurons in the NAc. These cells are involved in reward processing and are sensitive to the effects of stress (*19*–*22*). However, the specific effects of manipulating *Htra1* in astrocytes on NAc MSNs have not been studied. Therefore, we virally manipulated *Htra1* expression in NAc astrocytes and examined its effects on D1- and D2-MSN activity following exposure to ST-CVS using ex-vivo slice electrophysiology in male and female D2-GFP mice (Fig. 3A). This approach enabled us to record from both MSN populations proximate to our AAV5-GfaABC1D-*Htra1* OE and KD within the same animal. Strikingly, in males, *Htra1* KD combined with ST-CVS induced hyperexcitability in both D1- and D2-MSNs (Fig. 3B,C) compared to all other groups. Conversely, in females, *Htra1* OE combined with ST-CVS induced hyperexcitability exclusively in D2-MSNs (Fig. 3D,E) compared to all other groups.

**Fig. 3.**
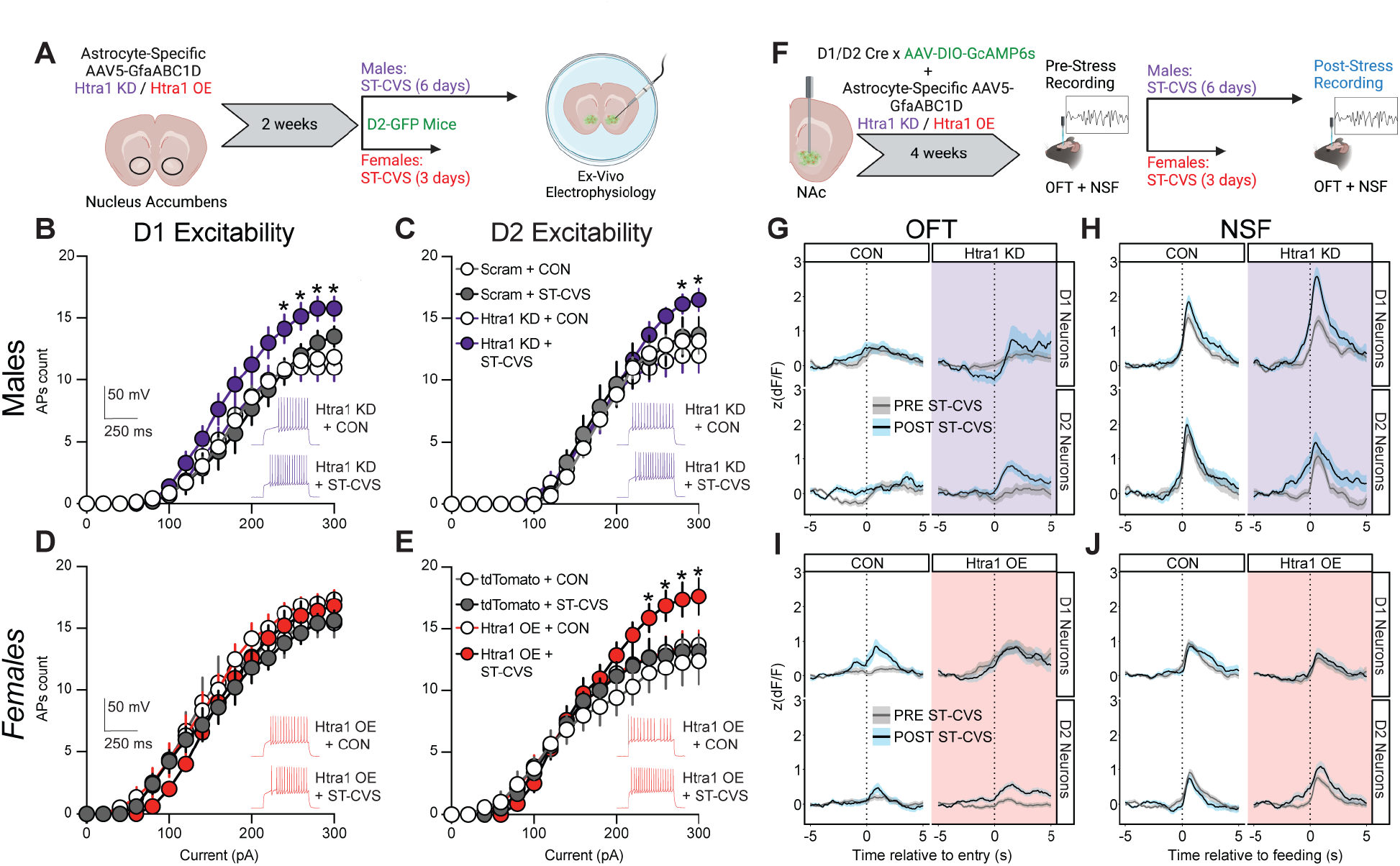
Astrocyte-specific regulation of *Htra1* in the NAc alters D1 and D2 MSN activity in a sex-specific manner. (**A**) Schematic outlining experimental approach of ex-vivo electrophysiological recordings in D2-GFP mice. In males, *Htra1* KD + ST-CVS produced a significant increase in (**B**) D1 MSN and (**C**) D2 MSN activity compared to all other groups (Three-way ANOVA: virus F_1,19.26_ = 0.0008 *p =* 0.97839, stress F_1,19.26_ = 0.8834 *p =* 0.35891, current F_1,804.01_ = 3788.1 *p* < 0.0001, virus x stress x current F_1,804.01_ = 15.682 p < 0.0001; followed by FDR corrected post-hoc tests). In contrast, in females, *Htra1* OE + ST-CVS produced no change in (**D**) D1 MSN excitability, but significantly increased excitability in (**E**) D2 MSNs (Three-way ANOVA: virus F_1,23.24_ = 2.4165 *p =* 0.1335, stress F_1,23.24_ = 2.9477 *p =* 0.0993, current F_1,806.01_ = 3787.7 *p* < 0.0001, virus x stress x current F_1,806.01_ = 4.9010 p = 0.0271; followed by FDR corrected post-hoc tests). (**F**) Schematic outlining experimental approach of in-vivo calcium imaging using fiber photometry recordings during behavior in D1/D2-Cre mice. (**G**) Average GCaMP6s signal in D1-Cre (top) or D2-Cre (bottom) male mice during entries to the center of the OFT before ST-CVS (grey) versus after ST-CVS (blue). (**H**) Average GCaMP6s signal in D1-Cre (top) or D2-Cre (bottom) male mice during feeding in the NSF before ST-CVS (grey) versus after ST-CVS (blue). (**I**) Average GCaMP6s signal in D1-Cre (top) or D2-Cre (bottom) female mice during entries to the center of the OFT before ST-CVS (grey) versus after ST-CVS (blue). (**J**) Average GCaMP6s signal in D1-Cre (top) or D2-Cre (bottom) female mice during feeding in the NSF before ST-CVS (grey) versus after ST-CVS (blue). Only traces when the mouse spent at least 1 second in the center of the OFT or feeding in the NSF were used for averaging. \**p*<0.05 compared to respective CON group; Abbreviations: KD, knockdown; MSN, medium spiny neuron; OE, overexpression; CON, control; CVS, chronic variable stress; ST-CVS, subthreshold CVS; OFT, open field test; NSF, novelty suppressed feeding.

It is important to note that ex-vivo electrophysiology findings may not always translate directly into *in vivo* changes. Therefore, to further elucidate the functional consequences of astrocytic *Htra1* manipulation, we evaluated D1- and D2-specific MSN activity in NAc *in vivo* using fiber photometry in awake, behaving mice (Fig. 3F & Fig. S4). Consistent with the ex-vivo electrophysiological results, D1- or D2-specific *in vivo* GCaMP6s calcium imaging in freely-moving male D1-Cre or D2-Cre mice revealed a significant increase in both D1- and D2-MSN activity exclusively in the *Htra1* KD group following ST-CVS in both the OFT and NSF tasks (Fig. 3G,H). In contrast, in females, we observed a significant increase in D2-MSN activity exclusively in the *Htra1* OE group following exposure to ST-CVS in the OFT and NSF tasks (Fig. 3I,J). Overall, these findings underscore the opposite effects of *Htra1* manipulation on stress-induced MSN activity in NAc both in brain slices and *in vivo*.

### Chronic stress and astrocyte HTRA1 alter perineuronal net intensity in the NAc in a sex-specific manner

One potential mechanism by which astrocytic HTRA1 could control the activity of NAc MSNs is through the regulation of ECM remodeling (*12*). Although the ECM is present in all tissues of the body, the brain’s ECM is notably different in composition and structure, varying from a widespread, uniform, shapeless matrix to well-defined structures such as basement membranes and perineuronal nets (PNNs) (*23*). The brain’s ECM is composed of a complex network of proteoglycans, glycoproteins, and hyaluronic acid, which surrounds neurons and glial cells, with minimal fibrous proteins unlike other tissues (*24*). The ECM provides structural support, guides cell migration, promotes cell maturation, fosters cell survival, and acts as a master regulator of synaptic plasticity (*25, 26*). Notably, ECM molecules, secreted into the extracellular space by embedded cells, can interact with nerve and glial cells via receptors that respond to specific ECM components (*24*). This dynamic structure, constituting about 20% of the adult brain’s total volume, undergoes continuous remodeling, with ECM components being deposited, degraded, or otherwise modified throughout life (*27*). This has led to the concept of the “tetrapartite synapse,” which posits that brain plasticity is governed by neurons and glial cells as well as by the ECM (*28*). Indeed, ECM alterations are increasingly linked to depressive symptomatology induced by chronic stress and present in MDD (*29, 30*). Proteoglycans, particularly those of the hyalectan/lectican family, are key structural elements and play crucial roles in the composition and function of the brain’s ECM (*31*). Brevican (BCAN), the most abundant lectican in the adult brain, controls synaptic plasticity and neuronal function, and is subject to proteolytic processing (*32*–*35*). Given that Htra1 is a secreted protease known to cleave several ECM proteins, including those in the lectican family (*12*), we hypothesized that manipulating HTRA1 expression in astrocytes could influence the availability of these ECM proteins, thereby influencing MSN activity.

To test this hypothesis, we first assessed BCAN protein expression levels in the NAc from postmortem human brains of patients with MDD. We found that, like HTRA1 expression, BCAN expression is sexually dimorphic and oppositely regulated between males and females in the context of MDD. Specifically, males exhibit significantly lower levels of BCAN expression compared to females at baseline, while male patients with MDD display significantly higher BCAN expression than their control counterparts, with female MDD patients showing a significant reduction in BCAN expression compared to their respective controls (Fig. 4A). Furthermore, we observed an inverse correlation between BCAN protein expression and HTRA1 protein expression in humans (Fig. 4B), supporting the notion that astrocyte-derived HTRA1 levels mediate ECM remodeling events by degrading BCAN. Next, we evaluated BCAN protein expression levels in male and female mice exposed to CVS (Fig. 4C), again noting sexually dimorphic and oppositely regulated expression between males and females in the context of chronic stress. Additionally, consistent with our observations in humans, we found an inverse correlation between BCAN protein expression and HTRA1 protein expression in male and female mice regardless of stress exposure (Fig. 4D).

**Fig. 4.**
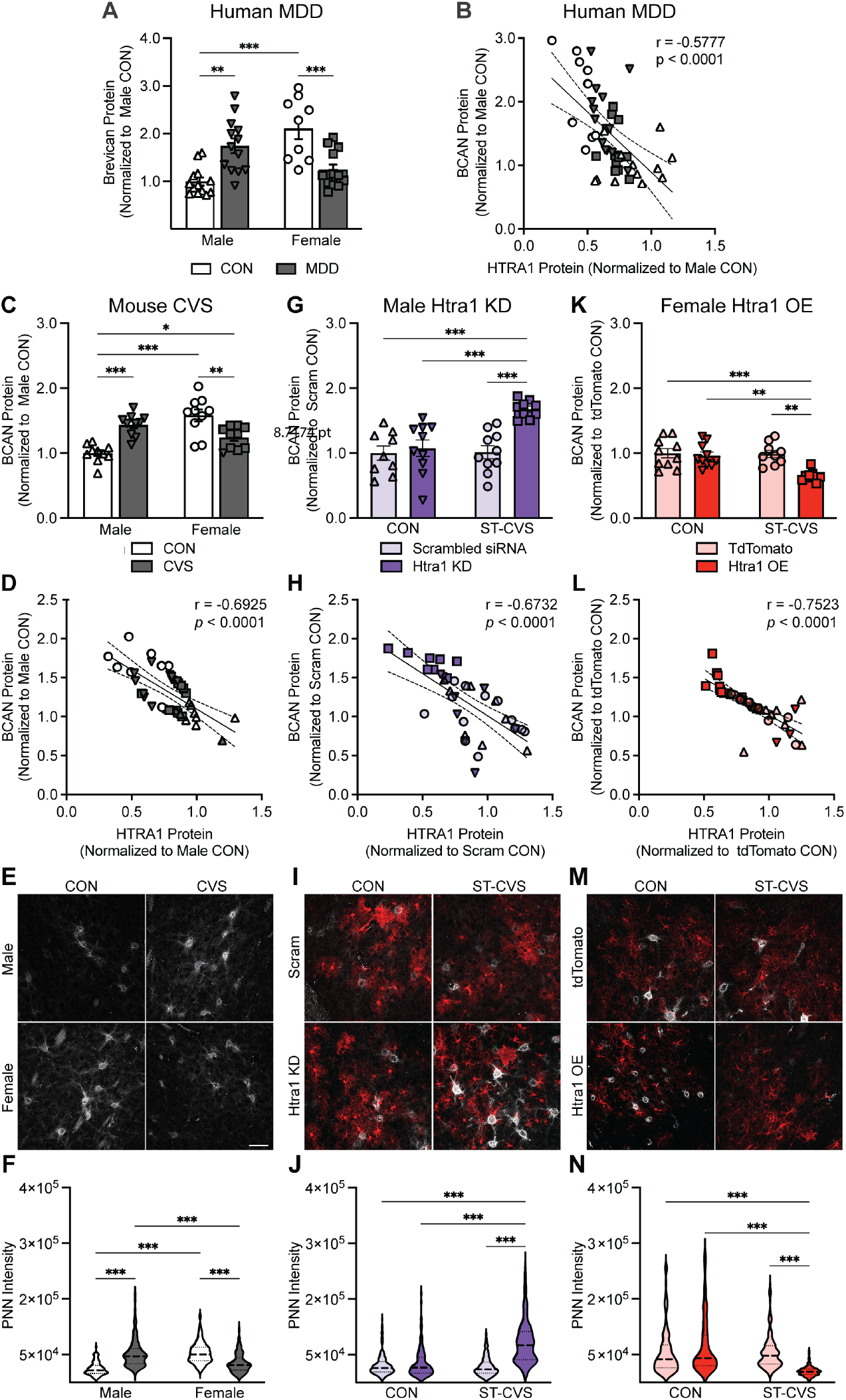
Stress and astrocyte-specific manipulation of *Htra1* expression in the NAc alter ECM-related organization in a sex-specific manner. (**A**) BCAN protein expression in the NAc is both sexually dimorphic as well as oppositely regulated in human MDD patients (Two-way ANOVA: sex F_1,44_ =4.848 *p* = 0.0330, MDD F_1,44_ = 0.1761 *p =* 0.6768, sex x MDD F_1,44_ = 33.67 *p* < 0.0001; followed by Tukey’s post-hoc tests). (**B**) Correlation between BCAN and HTRA1 protein expression in the NAc of human patients and controls. Line represents regression and its 95% confidence interval. Pearson’s correlation: *r* = -0.5777, *p* < 0.001. (**C**) BCAN protein expression in the NAc is both sexually dimorphic as well as oppositely regulated in mice exposed to CVS (Two-way ANOVA: sex F_1,36_ = 9.319 *p* = 0.0042, stress F_1,36_ = 0.6095 *p =* 0.4401, sex x stress F_1,36_ = 37.31 *p* < 0.0001; followed by Tukey’s post-hoc tests). (**D**) Correlation between BCAN and HTRA1 protein expression in the NAc of mice exposed to CVS. Line represents regression and its 95% confidence interval. Pearson’s correlation: *r* = -0.6925, *p* < 0.0001. (**E**) Representative confocal images of PNN staining in the NAc of CVS-exposed male and female mice. (**F**) PNN intensity in the NAc is both sexually dimorphic as well as oppositely regulated in mice exposed to CVS (Two-way ANOVA: sex F_1,15.9_ = 2.168 *p =* 0.1603, stress F_1,15.9_ = 0.9895 *p =* 0.3345, sex x stress F_1,15.9_ = 45.27 *p* < 0.0001; followed by Sidak post-hoc tests). (**G**) BCAN protein expression in the NAc of male mice is significantly increased following *Htra1* KD + ST-CVS (Two-way ANOVA: stress F_1,34_ = 13.47 *p =* 0.0008, virus F_1,34_ = 9.310 *p =* 0.0044, stress x virus F_1,34_ = 8.470 *p* = 0.0063; followed by Tukey’s post-hoc tests). (**H**) Correlation between BCAN and HTRA1 protein expression in the NAc of mice exposed to CVS. Line represents regression and its 95% confidence interval. Pearson’s correlation: *r* = -0.6732, *p* < 0.0001. (**I**) Representative confocal images of PNN staining in the NAc of male mice following *Htra1* KD in astrocytes. (**J**) PNN intensity in the NAc is significantly increased in males following *Htra1* KD + ST-CVS (Two-way ANOVA: virus F_1,15.6_ = 27.00 *p* < 0.0001, stress F_1,15.6_ = 11.91 *p =* 0.0033, sex x stress F_1,15.6_ = 20.02 *p* = 0.0004; followed by Sidak post-hoc tests).(**K**) BCAN protein expression in the NAc of female mice is significantly increased following *Htra1* OE + ST-CVS (Two-way ANOVA: stress F_1,34_ = 11.30 *p =* 0.0019, virus F_1,34_ = 8.266 *p =* 0.0069, stress x virus F_1,34_ = 7.316 *p* = 0.0106; followed by Tukey’s post-hoc tests). (**L**) Correlation between BCAN and HTRA1 protein expression in the NAc of mice exposed to CVS. Line represents regression and its 95% confidence interval. Pearson’s correlation: *r* = -0.7523, *p* < 0.0001. (**M**) Representative confocal images of PNN staining in the NAc of female mice following *Htra1* OE in astrocytes. (**N**) PNN intensity in the NAc is significantly decreased in females following *Htra1* OE + ST-CVS (Two-way ANOVA: virus F_1,14.9_ = 3.458 *p =* 0.0827, stress F_1,14.9_ = 10.86 *p =* 0.0049, virus x stress F_1,14.9_ = 8.038 *p* = 0.0125; followed by Sidak post-hoc tests).\**p* < 0.05; \*\**p* < 0.01; \*\*\**p* < 0.001. Abbreviations: MDD: Major Depressive Disorder; KD, knockdown; OE, overexpression; CON, control; CVS, chronic variable stress; ST-CVS, subthreshold CVS; PNN, perineuronal net.

To directly assess the impact of HTRA1 on BCAN levels, we examined the effects of astrocyte-specific KD or OE of *Htra1* on BCAN levels in NAc of male and female mice exposed to ST-CVS. We discovered that *Htra1* KD in males significantly increased BCAN protein levels, but only in the Htra1-KD + ST-CVS group compared to all other conditions (Fig. 5G). In contrast, we found that *Htra1* OE in females significantly reduced BCAN protein levels, again only in the Htra1-OE + ST-CVS group compared to all other conditions (Fig. 5K). Importantly, we observed an inverse correlation between HTRA1 and BCAN protein levels in both virally manipulated conditions (Fig. 5H,L). In light of these findings, it becomes imperative to directly study the influence of chronic stress and of astrocytic HTRA1 in ECM remodeling.

In the adult brain, ECM components are present primarily in intercellular spaces between neurons and glial cells (*36*). Lecticans, such as BCAN, interact with carbohydrate and protein ligands within the ECM, acting as linkers of these ECM molecules (*33*). Notably, they constitute primary components of perineuronal nets (PNNs), which are lattice-like formations of the ECM encapsulating neuronal cell body, proximal dendritic segments, axon initial segments, and presynaptic terminals (*37*). Recent focus on PNNs stems from the compelling notion that they restrict plasticity in adulthood but can be degraded to restore juvenile-like plasticity, facilitating axon sprouting and functional regeneration in damaged neurons (*38*). This underscores the pivotal roles of PNNs in synaptic stabilization, neuroprotection, and the regulation of ion homeostasis, particularly in neuronal populations characterized by heightened activity (*38*). Moreover, a growing number of studies investigating PNN contributions to neural and behavioral plasticity have utilized *Wisteria floribunda* agglutinin (WFA) fluorescence intensity as an indirect indicator of PNN maturity, where brighter staining signifies mature PNNs and dim staining indicates immaturity. Alterations in ECM, including changes in PNN deposition, have been observed following chronic stress exposure in rodents and implicated in human mood disorders (*39, 40*). Furthermore, astrocytes, as primary regulators of ECM dynamics in the brain, significantly contribute to ECM maintenance and remodeling, with adjacent astrocytic processes intricately intertwining with matrix material, contributing to the outer glial component of PNNs (*23, 41*). PNNs are present in the NAc (*42*), with recent findings suggesting BCAN’s crucial role in regulating PNN composition and dynamics in this region (*43*). Therefore, we assessed the effects of CVS exposure on PNN fluorescence intensity in the NAc (Fig. 4E), revealing sex-specific responses. Males exhibited lower baseline PNN fluorescence intensity compared to females, however, following CVS exposure, males showed a significant increase in PNN fluorescent intensity, while females displayed a significant reduction (Fig. 4F). These findings suggest a potential link between stress-induced alterations in PNN dynamics and the involvement of HTRA1. To explore this further, we manipulated *Htra1* expression directly in NAc astrocytes in male and female mice and assessed changes in PNN intensity following exposure to ST-CVS. In males, *Htra1* KD combined with ST-CVS induced a significant increase in PNN intensity compared to all other conditions (Fig. 4I,J). Conversely, in females, *Htra1* OE combined with ST-CVS induced a significant decrease in PNN intensity compared to all other conditions. Combined with the observed inverse correlation between HTRA1 and BCAN protein expression, we postulate that increased HTRA1 levels lead to reduced BCAN levels through degradation, resulting in diminished PNN intensity; conversely, decreased HTRA1 levels enable elevated BCAN levels, leading to heightened PNN intensity.

## Discussion

Our study delved into the transcriptional profiles of astrocyte-specific genes within the NAc of humans with MDD and of mice subjected to chronic stress. Among the identified genes, *HTRA1*, known for regulating the ECM, emerged as highly differentially regulated, displaying sexual dimorphism at baseline and opposite responses to depression or chronic stress between males versus females. We confirmed HTRA1’s predominant expression in astrocytes and showed that manipulating *Htra1* expression exclusively in NAc astrocytes bidirectionally regulated stress susceptibility in mice in a sex-specific manner. Moreover, our findings demonstrated that *Htra1* manipulation in NAc astrocytes modulated D1- and D2-MSN activity in brain slices and *in vivo* in a sex-specific manner, providing insight into the cellular mechanisms by which depression- and stress-associated changes in HTRA1 expression control stress susceptibility. Further investigations into the ECM revealed a direct relationship between HTRA1 and the proteoglycan BCAN. We observed sexually dimorphic and stress-responsive expression patterns of BCAN, inversely correlated with HTRA1 levels. Manipulating *Htra1* expression in NAc astrocytes controlled BCAN levels, providing a pathway through which HTRA1 regulates stress-induced ECM remodeling and consequent neuronal plasticity. We also revealed that chronic stress exposure altered PNN intensity in the NAc in a sex-specific manner. Strikingly, when we specifically manipulated *Htra1* in astrocytes of the NAc and exposed animals to ST-CVS, we observed a recapitulation of the changes in PNN intensity and BCAN protein expression that occur after CVS, once again demonstrating sex-specific effects.

Previous research has linked changes in PNN density or composition to several neuropsychiatric conditions including MDD. However, to date, most of these studies investigated PNNs in regions outside of the NAc, rarely accounting for sex. For example, studies in rodents have reported increased PNN deposition in the medial prefrontal cortex (PFC) associated with impaired cognitive processing and synaptic plasticity induced by chronic stress exposure, which can be ameliorated by PNN digestion, suggesting a link between PNN levels and mood (*40*). Moreover, rodents exposed chronically to either corticosterone or stress display heightened PNN deposition around fast spiking interneurons and elevated BCAN levels in hippocampus (*44, 45*). While studies on stress-induced changes in PNNs in humans are limited, postmortem samples from patients with bipolar disorder indicate alterations in PNN sulfation affecting proteolysis resistance in the amygdala (*46*). A history of child abuse is associated with increased densities and morphological complexity of PNNs in the ventromedial PFC (BA11/12), suggesting potential enhanced recruitment and maturation of PNNs (*47*). These observations point to the possibility that reversing PNN alterations may improve the associated behavioral abnormalities. Indeed, antidepressants like venlafaxine and ketamine have been found to reduce PNN levels, with implications for excitatory-inhibitory (E-I) balance, further supporting their role in mood regulation (*48, 49*). It is worth mentioning that excessive impairment of PNNs may also have adverse effects, particularly when considering our findings of contrasting changes in PNNs between males and females following exposure to chronic stress. For instance, mice lacking the PNN component neurocan exhibit behaviors resembling mania (*50*).

While PNNs are thought to predominantly enwrap parvalbumin (PV) inhibitory neurons in most brain areas (*51*), they surround excitatory neurons in the CA2 subfield of hippocampus (*52, 53*) as well as both PV^+^ and PV^-^ interneurons in the NAc (*54*). We confirmed this observation, finding that WFA^+^ PNNs in the NAc surround cells that are NeuN^+^ but DARP32^-^ (DARPP32 is a marker of D1- and D2-MSNs), indicating that they specifically surround interneurons and not MSNs in this region (Fig. S5). Although the intricacies of PNN effects on individual cells are complex, observations regarding neuronal population activity indicate that altered PNN deposition affects the balance between excitatory and inhibitory signaling (*40*). This is further supported by a study that investigated the impact of ECM molecule deficiencies on PNN formation, synaptic balance, and neuronal network activity, revealing altered synaptic composition, enhanced network activity, reduced PNN complexity, and differential gene expression in hippocampus of a quadruple knockout mouse model (tenascin-C^-^, tenascin-R^-^, BCAN^-^, and neurocan^-^) (*55*).

Our findings suggest a potential mechanism whereby astrocyte-derived HTRA1 influences ECM remodeling, particularly of PNNs located on interneurons within the NAc, influencing their E-I balance and ultimately affecting MSN activity. These results provide compelling evidence for the intricate interplay between ECM remodeling, astrocyte activity, and stress-response mechanisms in the modulation of neural circuitry underlying mood disorders that is impacted in a sex-specific manner. Further research is needed to understand the mechanisms underlying the opposite effects of depression and stress on HTRA1 and its downstream consequences in the NAc of males versus females as well as the opposite influence of this signaling pathway on stress-related behaviors between sexes. In conclusion, our study reveals a pivotal role of NAc astroglia and the ECM in mediating stress vulnerability that sets the stage for the development of sex-specific therapeutic interventions for mood disorders.

## Acknowledgments

The authors would like to thank Kyra Schmidt, Nathalia Pulido, Clementine Blaschke, Kinneret Rosen, and Ezekiell Mouzon for transgenic mouse breeding and genotyping.

## Author contributions

Conceptualization: EMP, EJN

Methodology: EMP

Investigation: EMP, TMG, AG, OKS, CJB, LFP, ATB, MS, RDC, MTR, AMC, LH, TM, YYZ, ZSL, FC, JBG, CT, OI, PJH

Formal analysis: EMP, AG

Visualization: EMP, AG

Funding acquisition: EMP, EJN

Supervision: EJN, CAB, SJR

Writing – original draft: EMP

Writing – review & editing: EMP, EJN

## Data and materials availability

All data are available in the main text or the supplementary materials. Custom R scripts and code utilized in this study, including for statistical analysis, are available upon request.

## Supplementary Materials

Materials and Methods

Supplementary Text

Figs. S1 to S5

Tables S1 References (58–66)

