## Supplemental Materials for "Sex-Specific Regulation of Stress Susceptibility by the Astrocytic Gene *Htra1*"

**The PDF file includes:**

Materials and Methods
Figs. S1 to S5
Table S1

### Materials and Methods

<u>Human brain postmortem</u>

Human postmortem brain tissue and RNA-seq data from MDD and control subjects were collected, analyzed, and reported as part of a published study (*58*). Briefly, brain tissue was obtained from the Douglas Bell Canada Brain Bank (Douglas Mental Health Institute, Verdun, Québec) and from the University of Texas Southwestern Medical Center Brain Bank. In total, analyses were performed on 89 samples including 25 male MDD, 25 female MDD, 17 male CTRL (healthy controls) and 22 female CTRL. Males and females were group-matched for age, pH, and BMI. Tissue was collected from six brain regions including the anterior insula (aINS), orbitofrontal cortex (OFC; BA11), cingulate gyrus 25 (BA25; cg25; vmPFC [ventromedial prefrontal cortex]), dorsolateral PFC (BA8/9; dlPFC), nucleus accumbens (NAc) and ventral subiculum (vSub). Detailed psychiatric history and socio-demographic information for all subjects included in the study have been previously reported (*56*, *58*). The study was approved by the research ethics boards of McGill University and UT Southwestern. Written informed consent was obtained from all participants. Here, we filtered these lists to identify astrocyte-specific genes (*59*) within the NAc with a nominal $P < 0.05$. NAc protein and cDNA from both cohorts was used to assess HTRA1 and brevican mRNA and protein levels.

<u>Pathway analysis</u>

Morpheus (https://software.broadinstitute.org/morpheus) was used to make heatmaps and perform hierarchical clustering analysis with default settings (one minus Pearson correlation and average linkage method). Gene expression data are plotted on the basis of log2 fold change. Criteria for presentation of GO terms included pathways containing a minimum of 5 genes and an adjusted *p*-value <0.05.

<u>Animals</u>

Animals were housed in the animal facilities at Mount Sinai. Male and female C57BL/6J mice (8-16 weeks old, 20-30 g, The Jackson Laboratory) were maintained on a 12:12 h light/dark cycle (07:00 lights on; 19:00 lights off) and were provided with food and water *ad libitum*. Transgenic mouse lines (D1-Cre: MGI:3836633, D2-Cre: MGI:3836635, and D2-eGFP: MGI:94924) were bred in-house on a C57BL/6J background. All animals were maintained according to the National Institutes of Health guidelines for Association for Assessment and Accreditation of Laboratory Animal Care accredited facilities. All experimental protocols were approved by the Institutional Animal Care and Use Committee at Mount Sinai.

<u>Viral reagents</u>

The plasmid for all AAV5-gfaABC1D constructs was obtained from Addgene (#44332). Additionally, the plasmid for *Htra1* overexpression was sourced from OriGene (Cat. No. MC201064), and the siRNA oligonucleotide set was acquired from Applied Biological Materials Inc (Cat. No. 240920940101). AAV vector production and purification were performed by Virovek (Hayward, California, USA) as previously described (*60*). Briefly, the AAV5-gfaABC1D-tdTomato-ScrambledRNA vector was generated in Sf9 cells through infection with rBV-inCap5-inRep-hr2 (Clone ID V295) and rBV-gfaABC1D-tdTomato-ScrambledRNA. Similarly, the AAV5-gfaABC1D-tdTomato-Htra1siRNA vector was produced using the same infection method with rBV-inCap5-inRep-hr2 (Clone ID V295) and rBV-gfaABC1D-tdTomato-

Htra1siRNA. AAV5-gfaABC1D-Htra1-IRES-tdTomato was generated in Sf9 cells through infection with rBV-inCap5-inRep-hr2 (Clone ID V295) and rBV-gfaABC1D-Htra1-IRES-tdTomato. Lastly, AAV5-gfaABC1D-tdTomato was produced in Sf9 cells through infection with rBV-inCap5-inRep-hr2 (Clone ID V295) and rBV-gfaABC1D-tdTomato. The final buffer for each virus was 1xPBS + 0.001% pluronic F-68. The purification process involved two rounds of CsCl ultracentrifugation, followed by CsCl removal through buffer exchange using 2 PD-10 desalting columns. The vectors were ultimately sterilized by filtration using 0.22 μm filters.

For calcium imaging, AAV9-CAG-Flex-GCaMP6s-WPRE-SV40 was obtained from Addgene (#100842). All viruses were used at $\pm 2.5 \times 10^{12}$ GC/mL, except AAV9-CAG-Flex-GCaMP6s-WPRE-SV40 at $\pm 5 \times 10^{12}$ GC/mL. Viruses were diluted to their appropriate titer using sterile PBS on the day of surgery.

RNA extraction and quantitative real-time PCR

Mouse brains were collected after cervical dislocation and followed by rapid bilateral NAc punch dissections from 1 mm-thick coronal brain sections using a 14G needle and frozen on dry ice. RNA extraction was performed using the RNeasy Micro Kit (Qiagen) following manufacturer instructions. RNA 260/280 ratios of the samples were confirmed using a NanoDrop Microvolume Spectrophotometer (ThermoFisher Scientific), and reverse transcription was achieved using the iScript cDNA Synthesis Kit (BioRad). Quantitative PCR using PowerUp SYBR Green (Applied Biosystems) was used to quantify cDNA using an Applied Biosystems QuantStudio 5 system. Each reaction was performed in quadtruplicate and relative expression was calculated relative to the geometric average of the control gene HPRT1 according to published methods (*61*). Sequences of the primers used can be found in Table S1 below.

**Table S1.** Primer sequences for qRT-PCR

| Gene | Species | Forward Sequence (5'–3') | Reverse Sequence (5'–3') |
|---|---|---|---|
| *HTRA1* | Human | TCCCAACAGTTTGCGCCATAA | CCGGCACCTCTCGTTTAGAAA |
| *Htra1* | Mouse | TAGCGACGCCAAGACCTACA | TGACGCAAACTGTTGGGATCT |
| *HPRT1* | Human | GACTAATTATGGACAGGACTGAACGTC | TCTCCTTCATCACATCTCGAGC |
| *Hprt1* | Mouse | GCAGTACAGCCCCAAAATGG | GGTCCTTTTCACCAGCAAGCT |

Protein extraction and Western blotting

Mouse brains were collected after cervical dislocation and followed by rapid bilateral NAc punch dissections from 1 mm-thick coronal brain sections using a 14G needle and immediately frozen on dry ice. For whole tissue extracts, frozen NAc samples were homogenized and then incubated for 30 min with agitation in 200 μL of ice-cold RIPA buffer (10 mM Trizma Base, 150 mM NaCl, 1 mM EDTA, 0.1% SDS, 1% Triton-X-100, 1% sodium deoxycholate, pH 7.4, complemented with protease and serine/threonine and tyrosine phosphatase inhibitors), before 5 cycles of 20 s on/off sonication using a Bioruptor (Diagenode). Samples were centrifuged for 15 min at 14,000 g to pellet insoluble debris and lipids, and supernatant was transferred to new tubes. Protein concentration was quantified using a Pierce BCA Protein Assay Kit (ThermoFisher Scientific). SDS-PAGE protein separation and Western blotting were performed according to manufacturer instructions. Equal amounts of proteins were mixed with 3-mercaptoethanol-supplemented Laemmli buffer (BioRad), heated to 95°C for 5 min before being separated by SDS-PAGE with Criterion Precast Gels (4 –15% Tris-HCl; BioRad) and transferred onto Immun-Blot PVDF 0.2 µm (BioRad) membranes. Membranes were blocked in Tris-buffered saline containing 5% bovine serum albumin (Sigma) and 0.1% Tween-20 (Fisher Bioreagents) for 1 h at room temperature. Primary antibodies anti-Htra1 (#PA5-11412, Invitrogen), anti-brevican (#19017-1-AP, Proteintech), and anti-β-Actin (#D6A8, Cell Signaling Technology) were diluted 1:500 in blocking solution and incubated overnight at 4°C. After washing, membranes were incubated with anti-rabbit (#7074, Cell Signaling Technology) peroxidase-conjugated secondary antibodies diluted 1:50,000 in blocking solution for 2 h, washed thoroughly, and developed using SuperSignal West Dura Substrate (ThermoFisher Scientific). Quantification was performed by densitometry using Image Studio (LI-COR Biosciences). Protein levels were normalized to actin. Between primary antibodies, membranes were stripped using Restore Plus Stripping Buffer (ThermoFisher Scientific). Mouse and human samples were run on separate gels and later normalized to their respective controls before being pooled for analysis.

Stereotaxic surgeries

Mice were anesthetized with an intraperitoneal bolus of ketamine (100 mg/kg) and xylazine (10 mg/kg), then head-fixed in a stereotaxic apparatus (Kopf Instruments). Syringe needles (33G, Hamilton) were used to bilaterally infuse 1 µl of virus at a 0.1 µl/min flow rate, except for fiber photometry experiments where infusion was unilateral. Needles were kept in place for 10 minutes after injection before being retracted to allow for virus diffusion. Coordinates for NAc were as follows, from Bregma: AP + 1.6 mm, ML + 1.5 mm, DV – 4.4 mm, 10° angle. For fiber photometry, 400 µm-wide optical fibers (Doric, MFC_400/430-0.66_4.5mm_MF2.5_FLT) were unilaterally implanted at AP – 1.6 mm , ML + 0.5 mm, DV – 4.3 mm, 0° angle. Virus infusion and optical fiber placement was confirmed either by immunohistochemistry on fixed brains sections or by dissection of fresh tissue under fluorescent light.

Immunohistochemistry and imaging

Mice were transcardially perfused with a fixative solution containing 4% paraformaldehyde (PFA). Brains were post-fixed for 24 h in 4% PFA at 4°C. Sections of 40 µm thickness were cut in the coronal plane with a vibratome (Leica) and stored at -20°C in a cryoprotectant solution containing 30% ethylene glycol (v/v), 30% glycerol (v/v) and 0.1 M phosphate buffer. Coronal brain sections through the nucleus accumbens (NAc; 1.94 mm through 0.74 mm from Bregma; Paxinos and Watson, 2007) were used. Astrocyte nuclei were labeled using an anti-Sox9 pimary antibody (rabbit; 1:1000; ab185966, Abcam), neuronal nuclei were labeled using an anti-NeuN primary antibody (mouse; 1:500; ab104224, Abcam; or rabbit; 1:500; ABN78, Millipore), MSN's were labeled using an anti-DARPP-32 primary antibody (mouse; 1/2,500; (*62*)), and native tdTomato signals from AAV5-GfaABC1D viral injections were enhanced using an anti-RFP primary antibody (goat; 1:1000; 200-101-379, Rockland). Sections were finally incubated with secondary antibodies donkey anti-rabbit Alexa Fluor 488 (1:500; 711-545-152, Jackson Immunoresearch), donkey anti-mouse Alexa Fluor 647, (1:500; 715-606-151, Jackson Immunoresearch), donkey anti-mouse Alexa Fluor 594 (1:500; 715-585-151, Jackson Immunoresearch) and donkey anti-goat Rhodamine-conjugated (1:500; 705-296-147, Jackson Immunoresearch), counterstained with DAPI and mounted in ProLong Diamond Antifade Mountant (ThermoFisher Scientific). Confocal images (1024 × 1024 pixels, 16 bits pixel depth, pixel size: x = 1.14 µm, y = 1.14 µm, z = 2.6 µm) were acquired on a LSM 780 upright confocal microscope (Zeiss) using a Plan-Apochromat 20x/0.8 M27 or Plan-Apochromat 40x/1.4 Oil DIC M27 objective and Zeiss Zen 2012 (black edition) software.

WFA staining

WFA staining was performed by washing free-floating sections three times for 5 min in 1X-PBS-Tx 0.2%. Sections were then incubated in 0.3% H202 in 0.2% PBS-TX for 20 min and then washed three more times for 5 min each in 1X-PBS-Tx 0.2%. The tissue was blocked in 2% bovine albumin serum in 0.2% PBS-Tx for 1 h and was then incubated overnight for two nights at 4 C on a rocking table with biotinylated WFA (1:500, Vector Laboratories) in 1X-PBS containing 2% BSA. To enhance the Native tdTomato signals, an anti-RFP antibody (1:500, Rockland Immunochemicals) was added 24 h into the two-day incubation, and sections continued to incubate for 24 h. The tissue was washed three times for 5 min each time in 1X-PBS-Tx 0.2% and then incubated in secondary antibodies streptavidin Alexa-Fluor 647 (1:500, Invitrogen) and donkey anti-Rhodamine (1:500, Jackson ImmunoResearch) for 4 h at room temperature. Finally, sections were incubated for 10 minutes in CuSO4 and then mounted in ProLong Diamond Antifade Mountant (ThermoFisher Scientific).

PNN imaging

Tissue was imaged using a LSM 780 upright confocal microscope (Zeiss) with a Plan-Apochromat 40x/1.4 Oil DIC M27 objective and Zeiss Zen 2012 (black edition) software was used for image acquisition. To ensure a comprehensive view of PNNs, imaging through a z-stack is essential, as PNNs surround neurons externally. Single-plane imaging may introduce bias and limit data acquisition. Therefore, images were taken through a z-plane (20 um) within the NAc, containing 20 stacks (1 um/stack). Laser intensity, gain, offset, and pinhole settings were kept constant for all images. Coronal brain sections through the nucleus accumbens (NAc; 1.94 mm through 0.74 mm from bregma; Paxinos and Watson, 2007) were used.

WFA quantification

Quantification of fluorescence intensity of PNNs was obtained in ImageJ using the Pipsqueak AI macro from Rewire Neuro Inc. (*63*). Individual regions of interest (ROIs) for each PNN were automatically created and verified by an experimenter who was blinded to the experimental condition. For all analyses, 4–6 images per animal were quantified.

RNA fluorescence in situ hybridization (FISH)

Mice were transcardially perfused with a fixative solution containing 4% paraformaldehyde (PFA). Brains were post-fixed for 24 h in 4% PFA at 4°C. Sections of 30 μm thickness were cut in the coronal plane with a vibratome (Leica) and stored at −20°C in a cryoprotectant solution containing 30% ethylene glycol (v/v), 30% glycerol (v/v) and 0.1 M phosphate buffer. NAc slices were mounted on charged Superfrost Plus microscope slides (Fisher Scientific) and processed for RNA FISH using RNAscope Multiplex Fluorescent Reagent Kit v2 (ACD Bio) according to manufacturer instructions using mouse probes for *NeuN* (Mm-Rbfox3, #313311), *Sox9* (Mm-Sox9-C2, 401051-C2) and *Htra1* (Mm-Htra1-C3, #423711-C3) transcripts. Sections were counterstained with DAPI and mounted using ProLong Diamond Antifade Mountant (ThermoFisher Scientific). Confocal images (6–10 per animal, 1024 x 1024 pixels, 16 bits pixel depth) were acquired on a LSM 780 upright confocal microscope (Zeiss) using a Plan-Apochromat

40x/1.4 Oil DIC M27 objective and Zeiss Zen 2012 (black edition) software. A custom-made automated ImageJ (U.S. National Institutes of Health) pipeline was used to extract channel intensity in every nucleus identified on DAPI staining (Fig. S1). Individual *Htra1, Sox9, and NeuN* puncta were detected using ComDet v 0.5.4 (github.com/ekatrukha/ComDet). Analysis code is available upon request.

*Ex vivo* slice electrophysiology

After at least 2 weeks of recovery from AAV surgery, male and female D2-GFP mice were subjected to ST-CVS or no stress and then anesthetized using isoflurane. Brains were rapidly extracted, and coronal sections (250 µm) were prepared using a Compresstome (Precisionary Instruments) in cold (0–4°C) sucrose-based artificial cerebrospinal fluid (SB-aCSF) containing 87 mM NaCl, 2.5 mM KCl, 1.25 mM $NaH_2PO_4$, 4 mM $MgCl_2$, 23 mM $NaHCO_3$, 75 mM Sucrose, 25 mM Glucose. After recovery for 60 min at 32°C in oxygenated (95% $CO_2$ / 5% $O_2$) aCSF containing 130 mM NaCl, 2.5 mM KCl, 1.2 mM $NaH_2PO_4$, 2.4 mM $CaCl_2$, 1.2 mM $MgCl_2$, 23 mM $NaHCO_3$, 11 mM Glucose, slices were kept in the same medium at room temperature for the rest of the day and individually transferred to a recording chamber continuously perfused at 2–3 mL/min with oxygenated aCSF. Patch pipettes (4–6 MO) were pulled from thin wall borosilicate glass using a micropipette puller (Sutter Instruments) and filled with a K-Gluconate-based intra-pipette solution containing 116 mM KGlu, 20 mM HEPES, 0.5 mM EGTA, 6 mM KCl, 2 mM NaCl, 4 mM ATP, 0.3 mM GTP (pH 7.2). Cells were visualized using an upright microscope with an IR-DIC lens and illuminated with a white light source (Olympus for Scientifica), and fluorescence visualized through eGFP and mCherry bandpass filters upon LED illumination through the objective (p3000$^{ULTRA}$, CoolLed). Excitability was measured in current-clamp mode by injecting incremental steps of current (0–300 pA, +20 pA at each step). For recording of spontaneous excitatory postsynaptic currents (sEPSCs), NAc MSN neurons were recorded in voltage-clamp mode at −70mV and detected with a 8 pA threshold. Whole-cell recordings were performed using a patch-clamp amplifier (Axoclamp 200B, Molecular Devices) connected to a Digidata 1550 LowNoise acquisition system (Molecular Devices). Signals were low pass filtered (Bessel, 2 kHz) and collected at 10 kHz using the data acquisition software pClamp 11 (Molecular Devices). Electrophysiological recordings were extracted using Clampfit (Molecular Devices). All groups were counterbalanced by days of recording and all recordings were performed blind to experimental condition.

Fiber photometry recordings and analysis

To analyze bulk photometry signals from GCaMP6s during mouse behavior, the fiber photometry system was time-locked with the video-tracking system (Ethovision XT 11, Noldus) via transistor–transistor logic signals (TTLs). The FP3002 system (NeuroPhotometrics) was used for all GCaMP6s recordings. It was connected to optical fiber head implants using low-autofluorescence patchcords, and was used according to manufacturer's instructions and with the FP3002 Bonsai node. Fluorescence signals resulting from 470 nm and 415 nm excitation, interleaved in time, were sampled at 78 Hz total, *i.e.* at a 26 Hz effective sampling rate for each channel.

Post-acquisition analyses were performed using custom programs. Deinterleaved NeuroPhotometrics time series data was directly exported from Bonsai. To compare neuronal activity across animals and behavioral sessions, individual animal time-series data using custom R codes following the standard method detailed in (*64*) with minor modifications. Briefly, the 470 and 415 nm signals were first smoothed using a 4th-order 5 Hz lowpass Butterworth filter built using the *gsignal::butter* function. To remove the bleaching slope and low-frequency fluctuations, baseline correction was then performed by subtracting the baseline obtained by regressing each individual signal using the LOWESS smoother (*stats::lowess*) with default parameters from the smoothed 470 and 415 nm signals. Both the 470 and 415 signals were then standardized using a robust z-score (zF = (F – median(F))/mad(F)). The standardized 405/415 nm signal was then fitted to the standardized 470 nm signal using the robust regression function *MASS::rlm*, and normalized dF/F z(dF/F) was finally calculated as the difference between the 470 nm signal and the fitted 415 nm signal to remove motion artifacts and autofluorescence. To analyze time-locked neuronal activity in respect to behavior, the normalized z(dF/F) signal was extracted around the onset of the relevant behavior (defined as t = 0 s). Finally, signal changes were quantified for relevant time intervals as the corresponding areas under the curve, which were calculated with linear interpolation using the *MESS::auc* function. Average signal trace data are averaged with n = event.

Chronic variable stress (CVS) procedure

CVS was performed as previously described (*56*). Male and female mice were subjected to a 21-day protocol consisting of three distinct stressors. To prevent habituation to the stress, the stressors were alternated throughout the 21-day period. The stressors were administered in the following sequence: first, 100 random mild foot shocks at 0.45 mA for 1 hour (with 10 mice in each chamber); second, a 1-hour session of tail suspension stress; and third, a 1-hour period of restraint stress where the mice were placed inside a 50 mL Falcon tube within their home cage. This sequence of three stressors is then repeated for the subsequent 21-day period. During the stress period, mice were housed in groups of five per cage, but at the start of behavioral assessments, the mice were rehoused individually. Control groups consisting of unstressed mice housed together were used for both males and females. Similar to the stressed mice, the control mice were also rehoused individually at the start of the behavioral experiments.

Subthreshold CVS (ST-CVS) procedure

A subthreshold version of CVS was used to study stress susceptibility; this duration of stress by itself is insufficient to induce stress-related behavioral abnormalities in males and females, but when combined with certain virally mediated modifications of gene expression can yield enhanced stress susceptibility (*65*). In males, the same stressors described above were repeated over a period of 6 d, whereas in females, stressors were repeated for 3 d since they are more sensitive to the effects of CVS (*57*). On day 7 in males and 4 in females, behavior was assessed in the splash test (ST), novelty suppressed feeding (NSF), sucrose preference (SP), and open field test (OFT) tests (in that order) as described below.

Splash test (ST)

The ST was performed as previously described (*56*). The test was performed under an incandescent red-light bulb (230 V, 15 W). Mice were habituated to the room for 1 h before testing. Mice were sprayed on the back with a 10% sucrose solution three times and then placed into an empty housing cage. Behavior was recorded for 5 min via videotape. The total amount of time grooming over the 5 min period was hand-scored by an observer blind to experimental conditions.

Sucrose preference (SP) test

Behavioral testing was conducted according to previously published protocols (*57*, *65*). Immediately after the splash test, mice were separated and individually housed. They were given two bottles filled with water for a 24-h habituation period. The following day, immediately after NSF testing, one of the two 50-ml bottles was replaced with a 2% sucrose bottle for 24 h. The two bottles were then weighed and their position was switched for an additional 24 h. The total duration of the test was 48 h. Sucrose was calculated by determining the percentage of total sucrose consumption divided by total liquid consumption (sucrose + water).

Novelty-suppressed feeding (NSF) test

The NSF test was performed as previously described (*56*). Mice were food-restricted for 24 h before testing occurs. Mice were then placed in the corner of a novel, black open-field arena (44 × 44 cm) covered in a different type of saw-dust bedding and where a single pellet of chow food was placed in the center of the arena. Testing conditions occurred under red-light conditions (10 lux) in a room isolated from external sound sources. The NSF boxes were thoroughly hand cleaned between mice with an odorless 30% ethanol cleaning solution, the bedding replaced, and a new food pellet was used for each mouse. Latency to feed was hand-scored by an experimenter blind to the experimental groups as the start of the first bout of uninterrupted feeding that was longer than 5 s.

Open field test (OFT)

Mice were placed in the open field arena (44 × 44 cm) for 10 min to compare the distance traveled and time spent in the peripheral zone compared to the center zone. Testing conditions occurred under red-light conditions (10 lux) in a room isolated from external sound sources. The OFT apparatus was thoroughly hand cleaned between mice with an odorless 30% ethanol cleaning solution. The mouse's activity – distance, velocity, and time spent in specific open field areas – was recorded using a video tracking system (Ethovision) set to localize the mouse center point at each time of the trial.

Statistics

All data are presented as mean ± S.E.M. Statistical analyses for were mainly conducted using Graphpad Prism 10.0, with a significance threshold set at $\alpha<0.05$ for all experiments. Statistical differences between two groups were analyzed with Student's t-tests with two-tailed analysis. Statistical differences between more than two groups were assessed using one-way or two-way ANOVAs, followed by Tukey's multiple comparison tests where applicable. The ROUT method was used to screen for outliers. Although specific statistical methods were not employed to determine the sample sizes, the number of experimental subjects was similar to the sample sizes routinely utilized in our laboratory and in the field for similar experiments (*66*). The data exhibited normal distribution, and variance was comparable between groups, providing support for the use of parametric statistics.

Statistics for the ex vivo electrophysiological recordings, fiber photometry, and the PNN intensity analyses were performed in R v4.2.2 mostly relying on *stats* v4.0.2, *tidyverse* v1.3.1 and *lmerTest* v3.1-3 packages. In summary, complex multifactorial designs were analyzed using linear models computed with the *stats::lm* function for fixed effects-only models or *lmerTest::lmer* function for mixed effects models. Random effects (repeated measures and/or nested observations) were modeled as random intercept factors. Subsequent analysis of variance was performed using type III sums of squares with Kenward-Roger's approximation of degrees of freedom. Post-hoc testing was performed using the *emmeans* package and significance was adjusted using Sidak's or FDR correction. Bar and line graphs represent mean ± sem. Significance was set at $p < 0.05$.

Data availability

Unmodified Western blots scans and other supporting raw data are available from the corresponding author upon request.

Code availability

Scripts and code utilized in this study, including for statistical analysis, are available from the corresponding author upon request.

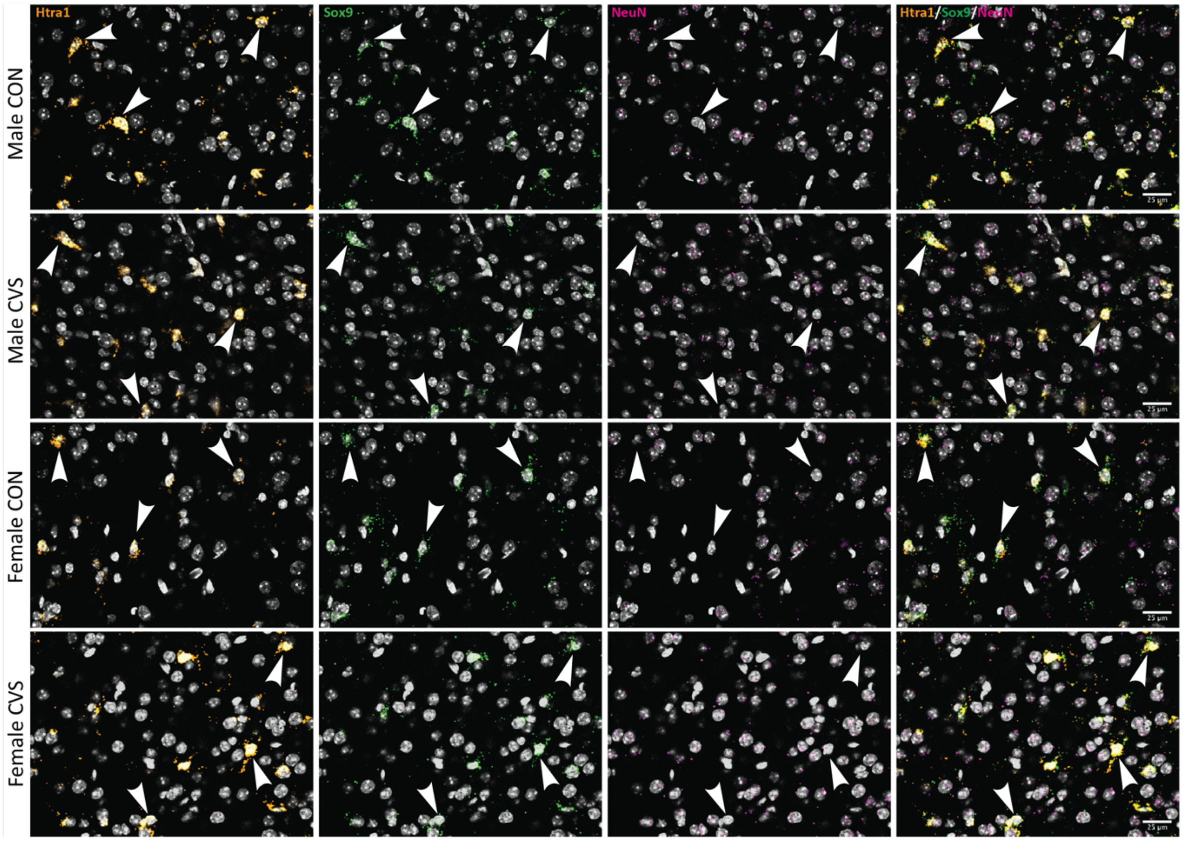


**Fig. S1. CVS-induced *Htra1* mRNA dysregulation is observed exclusively in astrocytes.** Representative confocal images (40x) from male and female mice exposed to CVS demonstrating that *Htra1* (orange) mRNA expression is significantly higher in astrocytes (green) compared to neurons (magenta) within the NAc of male and female mice regardless of stress exposure. Importantly, the previously observed sex- and stress-specific patterns of *Htra1* mRNA expression are exclusively observed in astrocytes. Arrows denote *Htra1*+ cells, and DAPI staining is in grey; Scale bars = 25 µm.

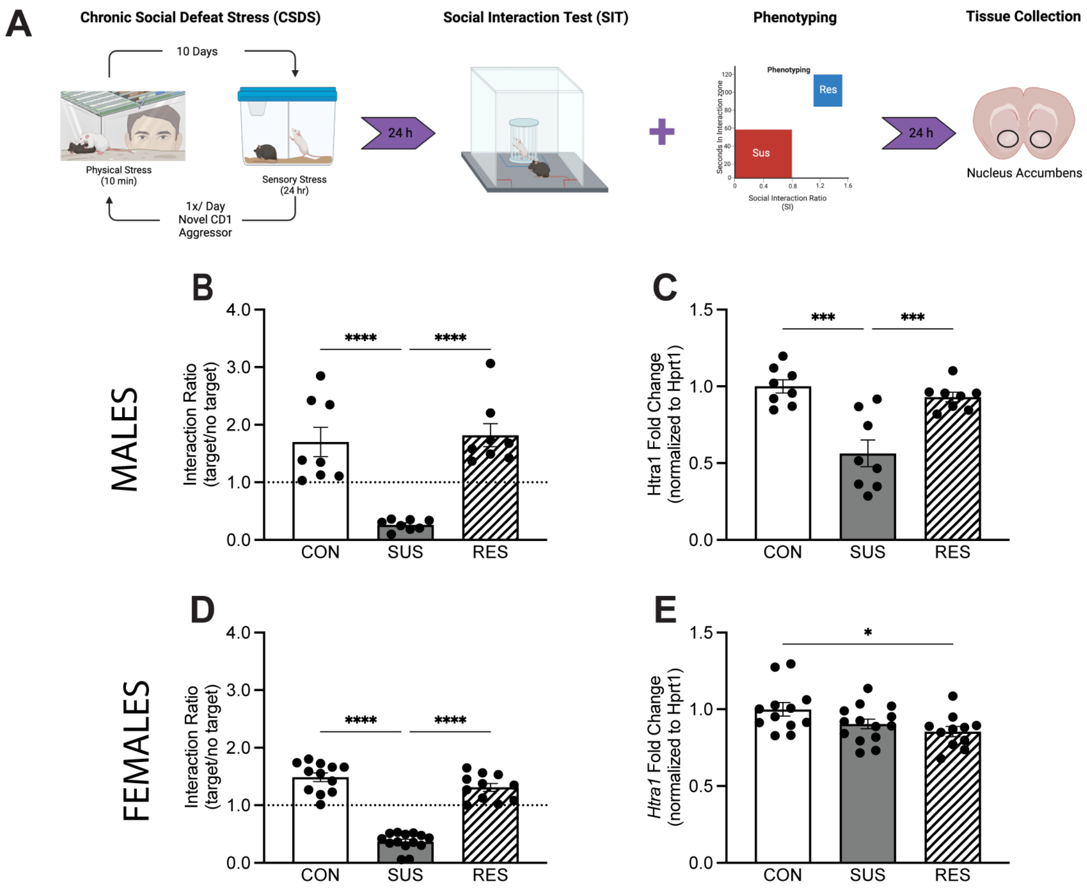


**Fig. S2. *Htra1* mRNA expression within the NAc is differentially regulated by CSDS in a sex-specific manner.** (**A**) Schematic depicting experimental design of CSDS exposure, behavioral phenotyping, tissue collection, and analysis. (**B**) In males, CSDS significantly reduces interaction ratio in susceptible (SUS) but not resilient (RES) mice compared to controls (CONs) (One-way ANOVA: $F_{2,21} = 21.41$ $p < 0.001$). (**C**) *Htra1* mRNA expression is significantly reduced in SUS male mice compared to CON and RES (One-way ANOVA: $F_{2,21} = 15.75$ $p < 0.001$) (**D**) In females, CSDS significantly reduces interaction ratio in SUS but not RES mice compared to CON (One-way ANOVA: $F_{2,34} = 103.6$ $p < 0.001$). (**E**) *Htra1* mRNA expression is significantly reduced in RES female mice compared to CON (One-way ANOVA: $F_{2,34} = 3.850$ $p < 0.05$). Abbreviations: CSDS, chronic social defeat stress; CON, control; SUS, susceptible; RES, resilient; NAc, nucleus accumbens.

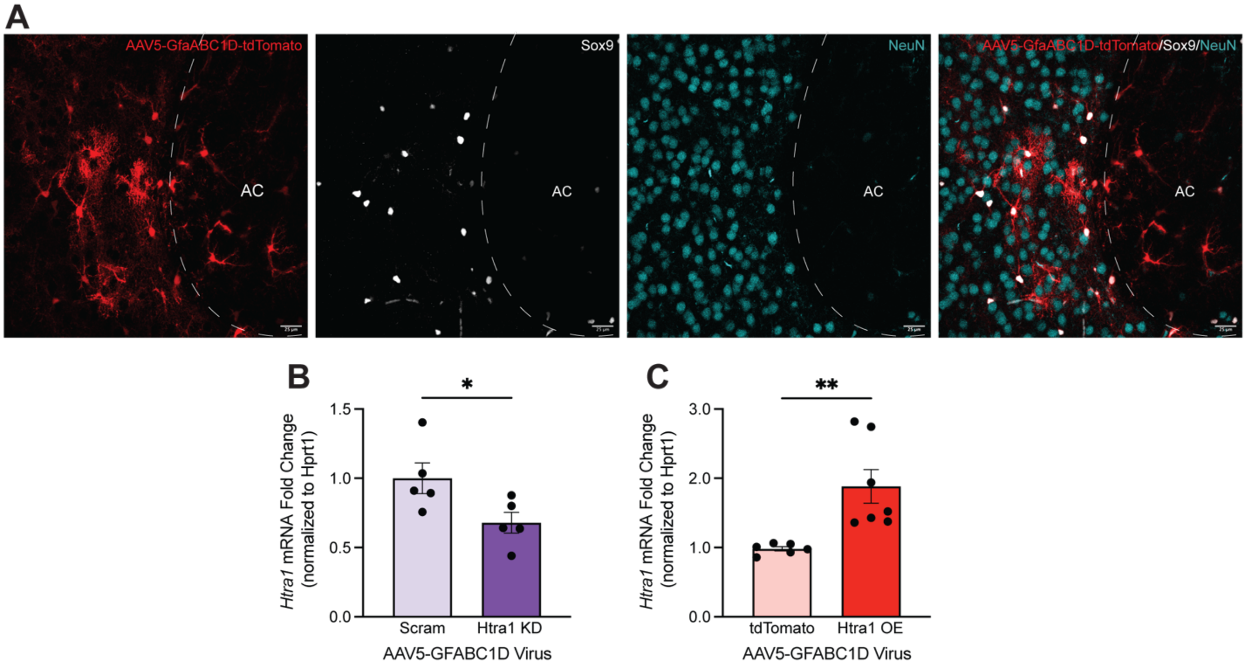


**Fig. S3. The AAV5-GfaABC1D vector exclusively infects astrocytes in mouse NAc**. (**A**) Representative confocal images (40x) of AAV5-GfaABC1D-GFP expression in the NAc alongside NeuN and Sox9, showing that all tdTomato+ cells colocalized with Sox9, with no overlap between NeuN+ and tdTomato+ cells. (**B**) AAV5-GfaABC1D-Htra1-siRNA significantly reduces *Htra1* mRNA expression compared to AAV5-GfaABC1D-Scram ($t_8$=2.436, p <0.05). (**C**) AAV5-GfaABC1D-Htra1 significantly increases *Htra1* mRNA expression compared to AAV5-GfaABC1D-tdTomato ($t_{17}$=3.150, p < 0.01). **p*<0.05; ***p*<0.01. Abbreviations: AC, anterior commissure; NAc, nucleus accumbens; KD, knockdown; OE, overexpression. Scale bars = 25 μm.

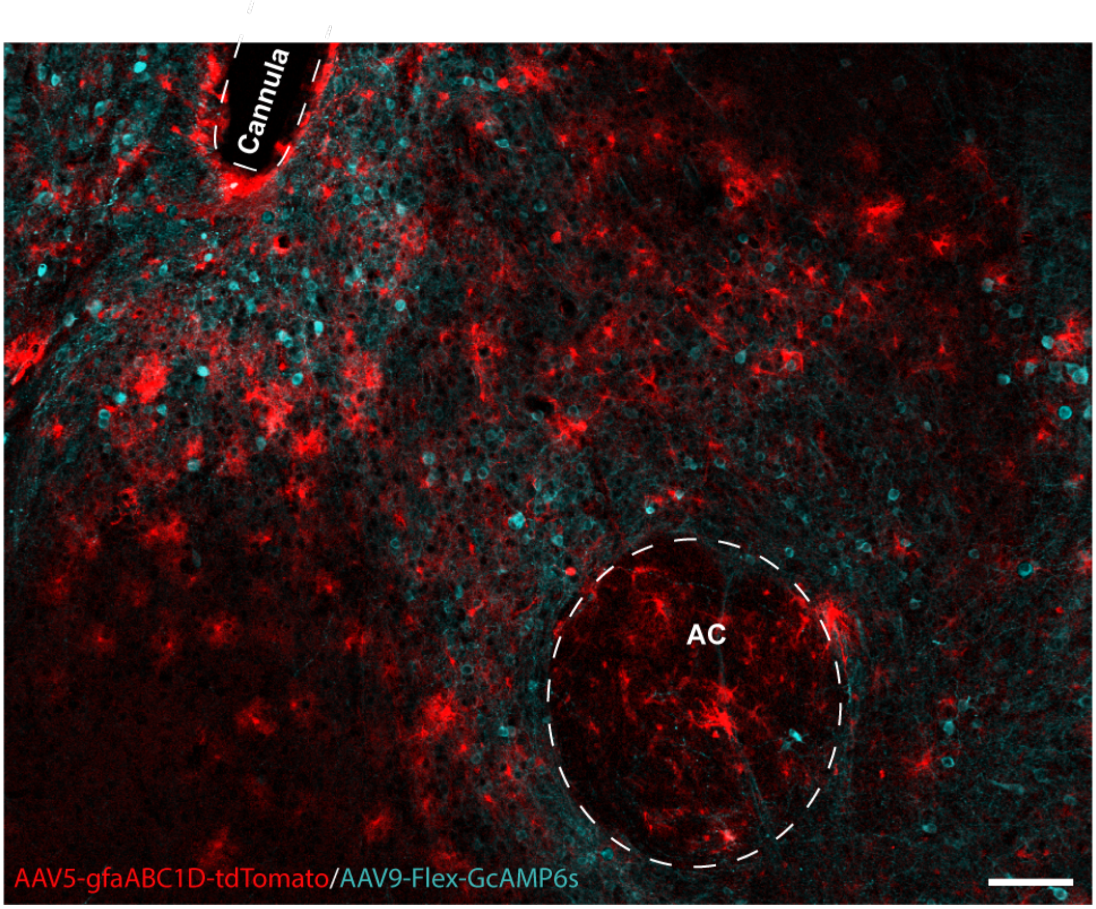


**Fig. S4. Representative confocal image (20x) of AAV5-gfaABC1D-tdTomato and AAV9-Flex-GCaMP6s expression in the NAc of a D1-Cre male mouse.** Dotted line depicts placement of a fiber optic cannula. Abbreviations: AC, anterior commissure; NAc, nucleus accumbens. Scale bar = 100 µm.

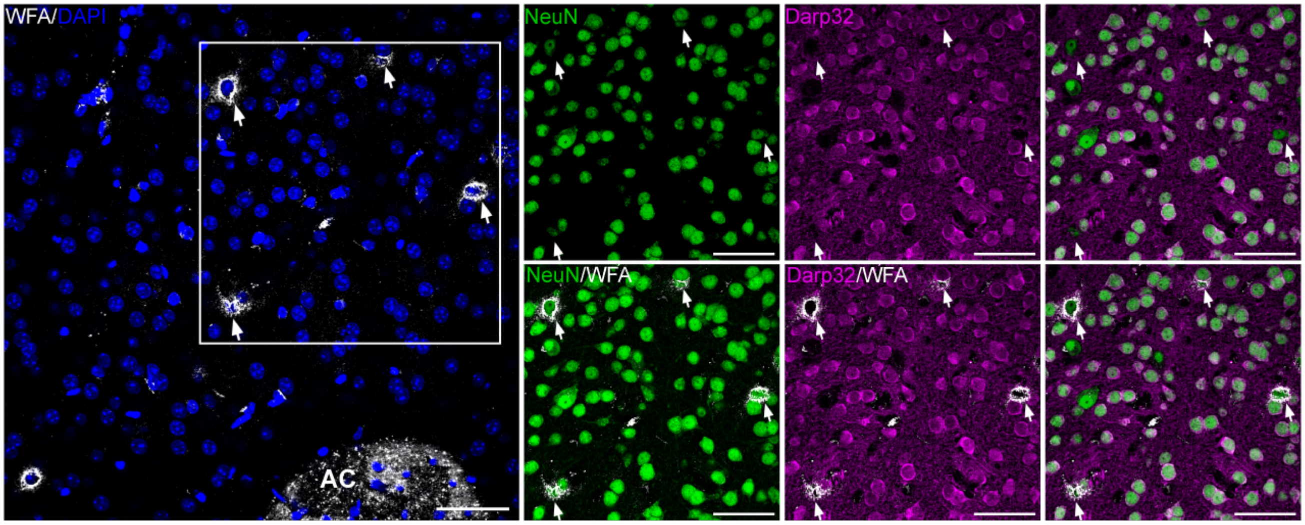


**Fig. S5. PNNs are located around interneurons but not MSNs in the NAc.** Representative confocal images (40x) of WFA staining (white) in the NAc alongside DAPI (blue)**,** NeuN (green), and DARPP32 (magenta), showing that WFA+ PNNs are NeuN+ but DARPP32-, indicating that they are located around interneurons rather than MSNs (denoted by the white arrows). Abbreviations: WFA, Wisteria floribunda agglutinin; PNNs, perineuronal nets; MSNs, medium spiny neurons. Scale bars = 50 µm.
